# Genomic stability and adaptation of beer brewing yeasts during serial repitching in the brewery

**DOI:** 10.1101/2020.06.26.166157

**Authors:** Christopher R. L. Large, Noah Hanson, Andreas Tsouris, Omar Abou Saada, Jirasin Koonthongkaew, Yoichi Toyokawa, Tom Schmidlin, Daniela A. Moreno-Habel, Hal McConnellogue, Richard Preiss, Hiroshi Takagi, Joseph Schacherer, Maitreya J. Dunham

## Abstract

Ale brewing yeast are the result of admixture between diverse strains of *Saccharomyces cerevisiae*, resulting in a heterozygous tetraploid that has since undergone numerous genomic rearrangements. As a result, comparisons between the genomes of modern related ale brewing strains show both extensive aneuploidy and mitotic recombination that has resulted in a loss of intragenomic diversity. Similar patterns of intraspecific admixture and subsequent selection for one haplotype have been seen in many domesticated crops, potentially reflecting a general pattern of domestication syndrome between these systems. We set out to explore the evolution of the ale brewing yeast, to understand both polyploid evolution and the process of domestication in the ecologically relevant environment of the brewery. Utilizing a common brewery practice known as ‘repitching’, in which yeasts are reused over multiple beer fermentations, we generated population time courses from multiple breweries utilizing similar strains of ale yeast. Applying whole-genome sequencing to the time courses, we have found that the same structural variations in the form of aneuploidy and mitotic recombination of particular chromosomes reproducibly rise to detectable frequency during adaptation to brewing conditions across multiple related strains in different breweries. Our results demonstrate that domestication of ale strains is an ongoing process and will likely continue to occur as modern brewing practices develop.

## Introduction

*Saccharomyces cerevisiae*, the common budding yeast, occupies a diverse number of environments, from an association with oak and fruit trees, to human-related industries such as baking and fermentation [1]. Modern efforts to characterize the diversity of *S. cerevisiae* through large whole-genome sequencing efforts have found a somewhat discrete population structure, in which strains isolated from a particular fermented beverage or geography are more closely related to other yeasts from that environment [2–5]. Due to a tight association with humans, the genomes of yeast are thought to have been shaped by both historical migrations of humans and the environment in which they are reared. One of the best characterized examples of this human-associated adaptation or domestication is the beer brewing yeasts, which are divided into three large clades across the family tree of yeasts. The largest division is split over species barriers between the *S. cerevisiae* ale yeast and the lager yeasts, which are hybrids between *S. cerevisiae* and *S. eubayanus* [6]. The ale yeasts are further divided into two large groups coined as Beer 1 and Beer 2, with several smaller mixed origin groups containing yeast from the bread, wine, and spirits industries. While the Beer 2 group consists primarily of diploid individuals used in traditional Belgian styles, the Beer 1 yeasts (herein called ale yeasts) consist of a diverse group of mostly tetraploid strains from Germany, Belgium, the UK, the USA, and Scandinavia [2,7]. The origin of the ale yeasts is hypothesized to come from a historical admixture between several *S. cerevisiae* subpopulations who have similar genomic signatures as the extant populations of European and Asian wine strains with some beer brewing strains that are no longer in existence [8]. The diversity and structure of these populations has allowed for extensive study of the specific molecular adaptations beer brewing yeasts have to their human-created environment, making them an excellent system in which to study the genetic basis of domestication.

Using a combination of genotype association and phenotyping in previous works, several specific genetic variations have been linked to traits that are either beneficial for the flavor of a particular beer or for growth in a beer brewing environment. First, in comparison to wild strains, the ale brewing yeasts lack the ability to produce the flavor compound 4-vinylguaiacol (4-VG), which is undesirable in certain beer styles, through the inactivation of two genes, *PAD1* and *FDC1* [9,10]. Interestingly, one lineage of ale yeasts which are specifically used for making wheat beers in which 4-VG is a desirable characteristic, retain functional alleles, highlighting the diversity and specialization of the domesticated strains [2,5]. Second, ale yeasts encode an expansion of genes involved in the uptake and breakdown of maltose and maltotriose, two uniquely important sugar sources for beer brewing [2,5]. Finally, both wine and ale brewing yeasts show evidence for loss of function alleles in *AQY1* and *AQY2* resulting in increased osmotolerance in high sugar content environments [5,11]. However, there is a functional allele of *AQY2* present in some of the ale beer strains, potentially indicating either a lack of selection for this allele or an environment-dependent selective benefit. As most of these putative adaptations have been identified because they are shared among most or all of the ale brewing yeasts (Beer 1), it is unclear to what extent there are additional genetic variations which are resultant from adaptive evolution or domestication within subsets of the beer brewing yeasts.

Furthermore, these single gene events have simplified the process of connecting them to potential phenotypes while other mutations prominent in the lineage are more difficult to interpret. Likely as a result of the reduced ability of ale brewing yeasts to complete meiosis and the increased mutation rate of both aneuploidy and mitotic recombination in tetraploids [12], tracts of homozygosity have been extensively observed in these yeasts. Similarly, in lager brewing yeasts, extensive aneuploidy and mitotic recombination between and within these two genomes have led to tracts of homozygosity favoring certain *S. cerevisiae* or *S. eubayanus* alleles [6,13–15]. Although it is unclear what the consequence of these intragenomic events are in ale and lager yeasts, previous works from our group and others have shown that loss of heterozygosity (LOH) caused by mitotic recombination in a previously heterozygous strain can lead to drastic fitness consequences on the time scale of short-term experimental evolution [16–20]. Furthermore, in other yeasts, such as *Candida albicans* and *Cryptococcus neoformans,* extensive aneuploidy and LOH can lead to diverse phenotypic outcomes such as increased drug resistance and competitive growth (reviewed in [21]). Finally, in mitotically dividing human cells, LOH of a non-functional tumor suppressor allele can lead to an increased risk of cancer progression including, among others, *BRCA1* mediated breast and ovarian cancer [22]. While LOH has been both observed frequently in ale yeasts and has been seen to have phenotypic consequences in other mitotic cell populations, it has yet to be linked conclusively to traits in ale brewing yeasts.

Additionally, ale yeasts’ reduced ability to effectively go through meiosis complicates traditional quantitative trait mapping approaches for interpreting genetic variation. Alternative approaches that do not rely on meiosis, such as experimental evolution and genetic screens, have provided valuable insights into adaptation generally, including the importance of specific mutations, copy number variation [23], and ploidy [24]. Therefore, we decided to study adaptation to the brewery by taking advantage of a form of experimental evolution already being conducted at breweries. Typically, professional brewers serially reuse populations of yeast to brew batches of beer in a practice known as repitching or backslopping to reduce the financial burden of constantly buying yeast and to give the yeast the opportunity to physiologically adapt to the brewery. The process begins when a brewery purchases a batch of a particular yeast strain at scale (population size of ~2 x 10^13^) from a propagation company. These yeasts have commonly been grown from a patch of yeast, derived from a clonal glycerol stock stored at −80°C. When a propagation company sends out these yeasts, they often grow the stock beyond the needs of a single brewery to meet the demand for a particular strain, meaning that there are many generations of yeast growth that occur before the yeast arrive in the brewery (minimum of ~50 yeast generations). Once the yeasts arrive at the brewery they are inoculated or ‘pitched’ into a cereal and grain derived beer medium or ‘wort’. After the completion of fermentation at 10-14 days, the yeasts will flocculate to the bottom of the fermenter and are then collected. The brewer will typically collect approximately a third of the yeast, avoiding the trub that is made up of hop and protein particulates, and repitch the yeast into the next fermentation vessel with fresh wort. The actual number of yeast cells that are transferred varies from brewery to brewery and is often modified to match the starting sugar content of the media and the current viability of the yeasts.

Brewers will often limit the number of times that yeast are repitched to ~8 reuses to reduce the possibility of a failed batch by contamination, physiological changes to viability and vitality [25–27], and taste profile changes due to altered physiology [28,29] or genetic mutation. However, there are conflicting results about how long repitching can be continued before beer brewing yeast will undergo a detectable genetic change by evolution. Early research on the genetic consistency of brewing yeasts found the possibility that genetic mutations can affect brewing-relevant characteristics by phenotyping clones isolated from populations of reused yeasts in a continuous use fermenter [30]. Further research by a separate group found changes in flocculation behavior in clone isolates from serially reused yeasts over several years, potentially due to a deletion mutation in a flocculation-related gene [31]. Additionally, one study looking at population samples from serial reuse of yeasts was not able to show any genetic mutation over 135 serial fermentations through the use of gel-electrophoresis based methods [32]. In contrast, some recent works on lager fermentations of buckwheat and quinoa beer have found potential alterations in chromosome length over the course of serial repitching [33]. Despite the recent evidence that genetic based changes in beer characteristics rarely occur over short-term repitching, there are striking phenotypic differences between brewing yeasts that are almost certainly caused by genetic variation. Matching these observations, it is a common practice among professional and home brewers to cultivate a yeast strain for an extended number of yeast pitches to generate a so-called ‘house strain’ with altered brewing characteristics indicating that genetic changes will likely occur over some relatively short time period in the brewery. However, the mutational basis, timing, and consequence of these changes has not been fully documented using modern high-throughput whole genome sequencing.

Herein we describe the effect of long-term repitching on brewing yeasts from samples collected in collaborations with multiple breweries across the United States and Canada who use an American brewing strain, serially-repitched for greater than 10 cycles. From these collaborations we either collected a time course across the pitches and sequenced several representative time points, or sequenced a starting and final population (Fig 1). Using a combination of short and long read (Illumina and Oxford Nanopore) sequencing, we found large-scale chromosomal rearrangements rising to a detectable frequency even within the first several generations of repitching. As well, we discovered a potential link between a specific mitotic recombination event and both growth phenotypes and flavor metabolite production.

**Fig 1.**
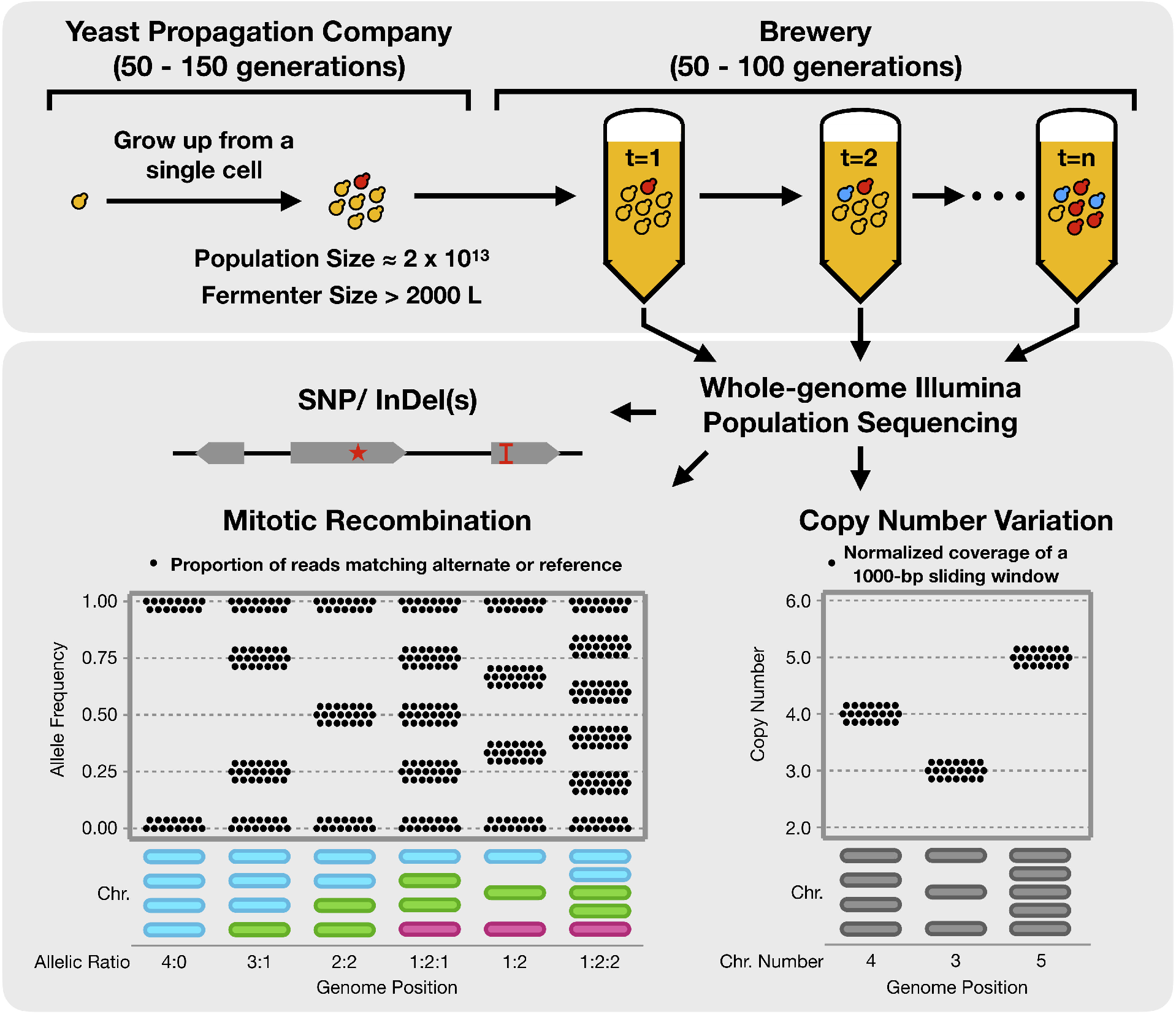
Research strategy to investigate a natural evolution experiment occurring in the brewery. Ale brewing yeast purchased from propagation companies at industrial scale and serially repitched for >15 beers are sequenced with whole-genome Illumina population sequencing to find signatures of evolution during brewery repitching. Mitotic recombination events are detected through allele frequency graphs (represented here), which denote every position of the genome as a point whose height is determined by the frequency of reads supporting either a reference or alternate allele. Copy number variation is detected and displayed as 1000-bp sliding windows, normalized by the average coverage across the genome.

## Results

We set out to tap into the natural evolution experiment occurring at modern breweries to test whether domestication is actively occurring in ale yeasts (Fig 1). We established collaborations with four breweries across the USA and Canada: Postdoc Brewing Co., Drake’s Brewing Co., Red Circle Brewing Co., and Elysian Brewing Co. All four breweries use a popular family of American yeast strains known as the ‘Chico’ yeasts and repitch for an extended number of cycles (>15), facilitating direct comparisons. Each brewery collected population samples of serially repitched yeast from independent beer lineages. For Postdoc Brewing Co. and Elysian Brewing Co. respectively we were able to collect two and three replicate beer lineages, plus one lineage each from the other two brewery partners. A complete record of the brewery populations is available in Table 1. Using whole-genome sequencing, we compared the starting genotype assessed from either a clone or population depending on availability with the last time point population sample for each beer lineage. For one replicate from Postdoc Brewing Co. we sequenced multiple population time points, and a number of representative clones isolated from the beginning and end time points for further experimental use. Given the time it takes to fully ferment a beer at an industrial scale, we estimate that the total time encapsulated in our experiments is on the order of four and a half years of yeast evolution.

**Table 1.** Record of strains from brewery collaborations

| Pop. Name | Brewery | Brewery ID | Strain | Rep. | Repitch Number | Variants Filtered By |
| --- | --- | --- | --- | --- | --- | --- |
| PDB1 | Postdoc Brewing Co. | PDB | Wyeast 1056 | 1 | 0 | N/A |
| PDB6 | Postdoc Brewing Co. | PDB | Wyeast 1056 | 1 | 6 | PDB1 |
| PDB15 | Postdoc Brewing Co. | PDB | Wyeast 1056 | 1 | 15 | PDB1 |
| PDB19 | Postdoc Brewing Co. | PDB | Wyeast 1056 | 1 | 19 | PDB1 |
| PDB26 | Postdoc Brewing Co. | PDB | Wyeast 1056 | 1 | 26 | PDB1 |
| PDB1 Rep. 2 | Postdoc Brewing Co. | PDB | Imperial A07 | 2 | 1 | N/A |

| PDB29 Rep. 2 | Postdoc Brewing Co. | PDB | Imperial A07 | 2 | 29 | PDB1 Rep. 2 |
| --- | --- | --- | --- | --- | --- | --- |
| E01 | Elysian Brewing Co. | Elysian. | BRY-96 | 1 | 0 | N/A |
| E03 | Elysian Brewing Co. | Elysian | BRY-96 | 1 | 15 | E01 |
| E08 | Elysian Brewing Co. | Elysian | BRY-96 | 2 | 15 | E01 |
| E09 | Elysian Brewing Co. | Elysian | BRY-96 | 2 | 17 | E01 |
| E05 | Elysian Brewing Co. | Elysian | BRY-96 | 3 | 1 | E07 |
| E07 | Elysian Brewing Co. | Elysian | BRY-96 | 3 | 0 | N/A |
| E10 | Elysian Brewing Co. | Elysian | BRY-96 | 3 | 14 | E07 |
| DK01 | Drake's Brewing Co. | Drakes | White Labs WLP001 | 1 | 24 | SRR7406282 |
| RCB01 | Red Circle Brewing Co. | Red Circle | Escarpment Cali Ale | 1 | 36 | SRR7406282 |
| Clone Name | Brewery | Brewery ID | Strain | Rep. | Repitch Number | Variants Filtered By |
| PDB1 c1 | Postdoc Brewing Co. | PDB | Wyeast 1056 | 1 | 1 | PDB1 |
| PDB1 c2 | Postdoc Brewing Co. | PDB | Wyeast 1056 | 1 | 1 | PDB1 |
| PDB26_c1 | Postdoc Brewing Co. | PDB | Wyeast 1056 | 1 | 26 | PDB1 |
| PDB26 c2 | Postdoc Brewing Co. | PDB | Wyeast 1056 | 1 | 26 | PDB1 |
| PDB26 c3 | Postdoc Brewing Co. | PDB | Wyeast 1056 | 1 | 26 | PDB1 |
| PDB26_c4 | Postdoc Brewing Co. | PDB | Wyeast 1056 | 1 | 26 | PDB1 |
| PDB26 c5 | Postdoc Brewing Co. | PDB | Wyeast 1056 | 1 | 26 | PDB1 |
| PDB26 c6 | Postdoc Brewing Co. | PDB | Wyeast 1056 | 1 | 26 | PDB1 |
| PDB26_c7 | Postdoc Brewing Co. | PDB | Wyeast 1056 | 1 | 26 | PDB1 |
| PDB26 c9 | Postdoc Brewing Co. | PDB | Wyeast 1056 | 1 | 26 | PDB1 |
| PDB26 c10 | Postdoc Brewing Co. | PDB | Wyeast 1056 | 1 | 26 | PDB1 |
| PDB26_c11 | Postdoc Brewing Co. | PDB | Wyeast 1056 | 1 | 26 | PDB1 |
| PDB26 c12 | Postdoc Brewing Co. | PDB | Wyeast 1056 | 1 | 26 | PDB1 |
| PDB26 c13 | Postdoc Brewing Co. | PDB | Wyeast 1056 | 1 | 26 | PDB1 |
| PDB26_c20 | Postdoc Brewing Co. | PDB | Wyeast 1056 | 1 | 26 | PDB1 |
| PDB26 c23 | Postdoc Brewing Co. | PDB | Wyeast 1056 | 1 | 26 | PDB1 |

As is common with most breweries, the same recipe was not used for each beer pitch, resulting in a potentially changing environment for the yeasts. Although these experiments are less controlled than traditional laboratory evolution experiments they provide a more realistic capture of the brewing environment. For the Postdoc Brewing Co. experiments, the order of different styles that the yeast went through is available in Supp. Table 1. Overall, the yeasts experienced an estimated final alcohol by volume (ABV) of around 5-6%. As well, for the Elysian samples, data collected at the brewery about the fermentation performance of each beer are available in Supp. Table 2 and show no strong deviation over the repitches.

Attempting to capture the full repertoire of genome variations that can occur and contribute to evolution, we investigated Single Nucleotide Polymorphisms (SNPs), Insertions and Deletions (InDels), copy number variations (CNVs), and changes in allele frequency resulting from mitotic recombination. First, in order to properly identify what variation occurred *de novo* during serial repitching, and how that variation relates to what occurs across the breweries, we established the relationship between the strains in our study cohort using phylogenetics.

### Relationship between strains

The history of the American brewing strains as told by brewers originates from just a handful of breweries. The Chico yeasts are specifically thought to originate from a ‘house-strain’ of the Sierra Nevada Brewing Company’s isolate of BRY-96, which is sold by the Siebel Institute. BRY-96 itself is thought to originate from P. Ballantine and Sons Brewing Company, which started in 1840 in Newark, New Jersey. The strain has since been distributed to a large number of breweries and yeast propagation companies. To provide a fuller picture of the genetic history of the American brewing yeasts, we collected not just the strains used by our brewery partners but also new clone samples of American brewing strains that are available for purchase and not believed to have been previously sequenced. In all, we sequenced 13 American brewing strains, and reanalyzed an additional 17 strains that had previously been sequenced using short-read sequencing (Supp. Table 3). Wanting to confirm the relationships between our study cohort, we applied phylogenetic inference on the strains. From their whole-genome sequence, we built a maximum likelihood tree based on the variation between these strains. However, as mentioned earlier, because there has been extensive mitotic recombination in these yeasts, we suspected that phylogenetic inference could be influenced by large blocks of shared, ancestral variation being lost. To avoid this issue, we filtered the American brewery strains variant calls by the most diverged American strain, BE051, to control for the potential loss of shared variation. As well, to encapsulate the polyploid nature of the beer strains, we encoded heterozygous variation in the genome sequences for phylogenetic inference (see methods for more details).

Matching with oral history, we found from our constructed phylogeny that Wyeast 1056 (Postdoc Brewing Co.), Imperial A07 (Postdoc Brewing Co.), White Labs WLP001 (Drake’s Brewing Co.) and Escarpment’s Cali Ale (Red Circle Brewing Co.), and other Chico yeasts are all closely related and form two large clades (Fig 2). As well, we found that the WLP001 and Wyeast 1056 clades are likely derived from BRY-96 (Elysian Brewing Co.), as there is only an 11 SNP difference between a reconstructed common ancestor of the two Chico strains and an isolate of what is thought to be the original BRY-96 (kindly donated by Lallemand Inc.). Additionally, from a sequenced isolate of a strain from P. Ballantine and Sons Brewing Company that was deposited in a strain repository in 1972 (NRRL Y-7408), we found that this strain groups outside of the rest of the American brewing strains, indicating that it is indeed a diverged American brewing strain. However, because large segments of variation are lost from NRRL Y-7408 that exist in the internal American brewing strains, we suspect that the Ballantine strain is not the literal genetic ancestor.

**Fig 2.**
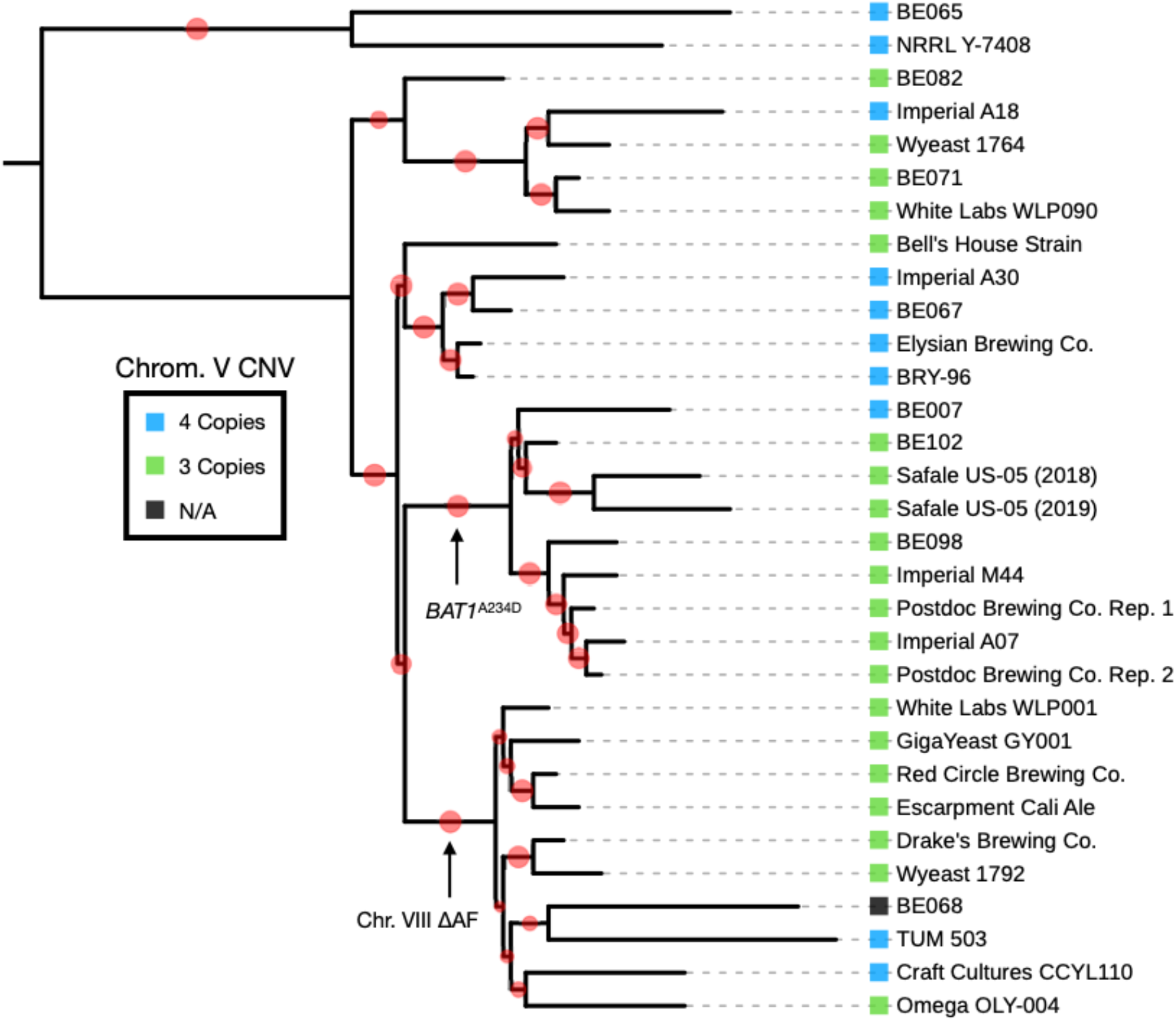
A maximum-likelihood phylogeny of the American brewing strains reveals several large clades. Specifically, two Chico yeast groups, and their presumed genetic ancestor, BRY-96 were found to group with other commercially available strains. The branch support bootstrap values are displayed in red on the adjoining branch, with smaller values corresponding to less support.

### *De novo* Single Nucleotide Polymorphisms, Insertions, and Deletions

With the ancestral strain sequences in hand, we next called *de novo* mutations that occurred during each repitching time course. Utilizing multiple SNP and InDel variant callers on the first replicate of the Postdoc Brewing Co. populations we did not find any *de novo* SNP or InDel that occurred during the course of the repitching experiment and reached a detectable frequency (estimated detection limit of ~2% of alternate reads). Using sequencing of clones isolated from the first time point to filter the variant calls from the populations, we found 11 mutations that were shared by all of the Postdoc Brewing Co. time points and had occurred in the population before entering the brewery based on the sequences from the clone isolates, the population from the second Postdoc Brewing Co. replicate, and the Imperial A07 clone isolate. Calculating the change in frequency of these mutations over the time course, we found that the only mutation that changed by more than a 1% increase in the population was a synonymous mutation in *PTC6* (which increased from 25.2% to 44.4% in the population, Supp. Table 4). While it is known that synonymous mutations can impact traits, it’s more likely that this is a passenger mutation, particularly since the mutation affects only one allele in a pentaploid region of the genome. We additionally observed a number of private SNPs and InDels within clones from both the first and last time points, with an average of 11.9 mutations per clone and a total of 177 unique mutations (Supp. Table 5 and Supp. Fig 1).

Expanding our analysis to the samples from the other collaborations, we found a total of 106 mutations, with an overall average of 15.1 mutations observed in each population (Supp. Table 5). Looking for evidence of adaptive evolution through convergence of mutations, we found that between experiments, there were 5 genes wherein multiple mutations were observed in the coding sequence between experiments (Table 2). We note that mutations in *UBP1,* which encodes a ubiquitin protease, were previously identified in experimental evolution of a lager strain [34], and mutations in *TFB1,* a nucleotide excision repair factor and subunit of TFIIH, were found in strains that had survived for two years in a sealed beer bottle [35]. However, in neither of these cases were phenotypic consequence proven. Further experiments recreating these mutations in clean genetic backgrounds will be necessary to determine their impact.

**Table 2.** The *de novo* mutations within the same genes between brewery populations

| Sample | Tool | Chromosome | Location | Ref | Alt | MutationType | Gene | Gene | Effect | FinalAF |
| --- | --- | --- | --- | --- | --- | --- | --- | --- | --- | --- |
| PDB29 Rep. 2 | freebayes | chrIV | 244970 | G | T | coding-nonsynonymous | YDL122W | UBP1 | D807Y | 0.103 |
| Red Circle Brewing | lofreq | chrIV | 244223 | G | T | coding-nonsynonymous | YDL122W | UBP1 | E558Stop | 0.040 |
| Elysian09 | lofreq | chrIV | 188256 | G | C | coding-nonsynonymous | YDL148C | NOP14 | Y777Stop | 0.101 |
| Elysian10 | lofreq | chrIV | 188256 | G | C | coding-nonsynonymous | YDL148C | NOP14 | Y777Stop | 0.129 |
| Elysian10 | lofreq | chrIV | 1085392 | C | T | coding-nonsynonymous | YDR311W | TFB1 | R110W | 0.108 |
| Drake's Brewing Co. | lofreq | chrIV | 1085083 | G | T | coding-nonsynonymous | YDR311W | TFB1 | A7S | 0.077 |
| PDB26 Rep. 1 | freebayes | chrXI | 245205 | C | G | coding-nonsynonymous | YKL104C | GFA1 | G57R | 0.077 |
| PDB29 Rep. 2 | freebayes | chrXI | 245205 | C | G | coding-nonsynonymous | YKL104C | GFA1 | G57R | 0.117 |
| Elysian03 | lofreq | chrXII | 905505 | G | A | coding-synonymous | YLR392C | ART10 | N267N | 0.176 |
| Elysian08 | lofreq | chrXII | 905987 | G | A | coding-nonsynonymous | YLR392C | ART10 | P107S | 0.049 |

### *De novo* chromosome copy number variation

We next investigated whether there were any large scale genomic changes by plotting the read coverage across the genome for the Postdoc Brewing Co. time course. In line with previous work, we observed that the ‘Chico’ yeasts are largely tetraploid, and have had several whole chromosome and segmental copy number changes (CNVs) that occurred at some point in its recent past (Fig 3A). We observed that during the Postdoc Brewing Co. time course there was a copy number chance of chromosome V which led to an increase from 3 to 4 chromosomal copies at a final estimated frequency in the population of 48.2% (Fig 3B). The second Postdoc Brewing experiment also replicated the increase in copy number on chromosome V (Fig 3C). Interestingly though, and unlike in the first Postdoc Brewing Co. replicate, the mutation entered the brewery at an estimated frequency of 27.6% and reached a frequency of 94.9% by the end of the experiment. For the population from Drake’s Brewing Co., we observed an estimated frequency of 27.9% for the chromosome V increase in copy number, indicating another replication of the same mutation. While no starting population was available for the sample, two separate sequences of WLP001 were uploaded two years apart by different groups and both shared 3 copies of chromosome V, indicating that the starting strain likely had 3 copies. Additionally, the strain from Red Circle Brewing Co., which is of similar origin, maintained 3 copies of chromosome V.

**Fig 3.**
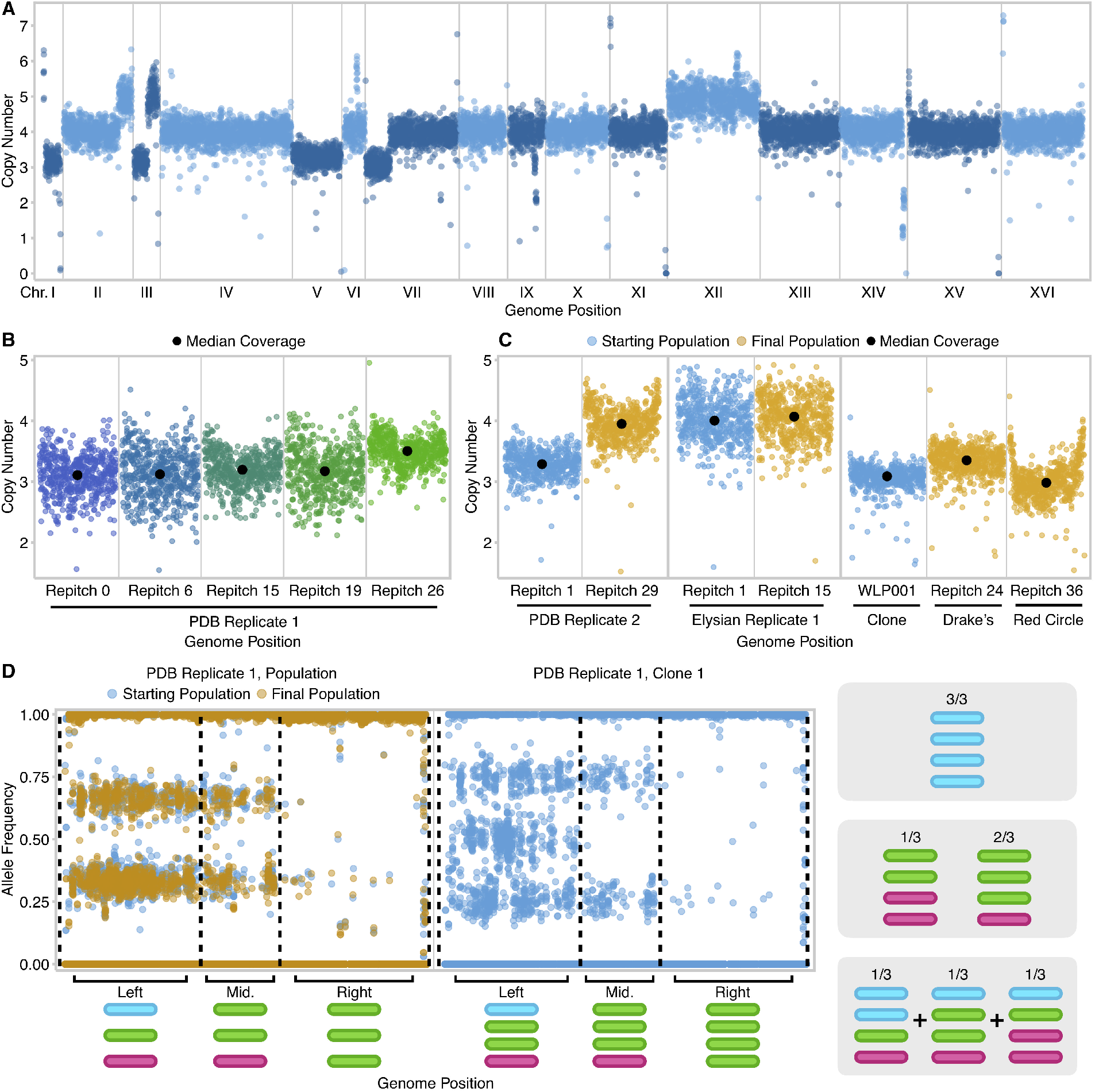
Copy number variation of chromosome V occurs multiple times between breweries and within the same brewery. (A) Whole-genome coverage of Imperial A07, highlighting the degree of chromosomal and sub-telomeric copy number alterations. (B) A time course of the Postdoc Brewing Co. replicate 1 is shown in 1000-bp coverage windows. A copy number change of chromosome V reaches 48.2% of the population by repitch 26. (C) The copy number increase of the second Postdoc Brewing Co. replicate population, starting at 27.6% of the population and reaching fixation by the 29th repitch. The strain BRY-96, which is used in Elysian Brewing Co. starts with a euploid copy number and remains constant during repitching. Drake’s Brewing Co., which is from WLP001, has an aneuploid lineage which reaches 94.9% of the population. The sample from Red Circle Brewing Co. showed an increase coverage near the telomeres across its genome, but this is likely a well-documented artifact [2] (D) Allele frequency of the Postdoc Brewing Co. replicate 1 population and a clone isolated from that population showing the number and pattern of haplotypes on chromosome V. The lack of a shift in allele frequency indicates that in the population, multiple lineages likely independently had different haplotypes amplified. The probability of any given haplotype being amplified is displayed on the right.

Wanting to determine whether the potential benefit of the aneuploidy was due to an increase in copy number of a particular haplotype or a restoration of a euploid copy number for dosage balance, we investigated whether one particular copy of chromosome V was recurrently amplified between populations. Our expectation is that gaining a chromosome copy will change the allele frequency of heterozygous variants by a change in the proportion of haplotypes. Through investigation of the direction that variants change allele frequency, we can determine which chromosome is amplified (See Fig 1 for allele frequency plot description). Therefore, we investigated whether the allelic ratio between haplotypes had changed by plotting the allele frequency of variants on chromosome V for the two Postdoc Brewing Co. and Drake’s Brewing Co. experiments. However, upon plotting the allele frequency from the first and last time points we found very little to no change had occurred despite the chromosome copy number change (Fig 3D and Supp. Fig 2).

From a clone isolated from the final population of the Postdoc Brewing Co., replicate 1 experiment that had an extra copy of chromosome V, we found that, in a clonal sample, as expected, the allele frequency does change and shows three large chromosomal regions with different allele frequency patterns. The clone helped show that the starting strain has three haplotypes on the left arm, two in the middle in a 2:1 ratio and is homozygous on the right arm. Given these patterns, we expect that depending on which chromosome was amplified, the allele frequency will shift according to the number of haplotypes (Left: 0.33/0.66 to 0.25/0.50/0.75; Center: 0.33/0.66 to 0.25/0.75 or 0.50; Right: No change, summarized on the right of Fig 3D). However, because the allele frequency pattern did not change in a significant manner, we instead concluded that there are likely multiple mutation events, each of which amplified a different chromosome V haplotype. These independent mutations have occurred in separate lineages that have risen in frequency with similar kinetics. Therefore, we suspect that the increase in copy number of chromosome V likely occurred multiple times in both Postdoc Brewing Co. replicates, indicating a haplotype independent fitness benefit.

The experiment(s) at Elysian Brewing Co. utilized BRY-96, which already contained 4 copies of chromosome V and did not show any additional evidence of aneuploidy. It is likely that the ancestral state of chromosome V is euploid based on the phylogenetic relationship between the American brewing strains (Fig 2). Since the chromosome loss event appears to have occurred multiple times in the ‘Chico’ phylogeny (Fig 2), it’s possible that this state could be selectively advantageous in certain environments. An alternative explanation is that when the common ancestor of WLP001 and Wyeast 1056 was clone isolated, the single-cell bottleneck fixed a deleterious mutation for growth, which was then reverted upon serial passaging in brewery conditions.

### *De novo* changes in heterozygosity

We next investigated whether there were copy number neutral changes in heterozygosity due to mitotic recombination by plotting the allele frequency of all positions in the genome. First, looking at the allele frequency of the Postdoc Brewing Co. populations over the sampled repitches, we observed a marked shift on the right arm of chromosome VIII starting at repitch number 15, angling towards an allele frequency of 0.50 (Fig 4A). Using the allele frequency of positions at the terminal end of the chromosome, we calculated that the allele frequency change reached a frequency of 43.8% by the end of the experiment. From clones isolated from the first replicate of the Postdoc Brewing Co. experiment, we identified that there were numerous, private breakpoints in each clone where the allele frequency changed from a haplotype ratio of 3:1 to 2:2 (Fig 4C; full list at Supp. Fig 3). We determined that the signal from the individual clone breakpoints stacked in the population data to create the angled pattern, with all events sharing a 2:2 ratio at the most distal segment of the chromosome. From these data, we concluded that there are two chromosomal haplotypes on the right arm of chromosome VIII in a 3:1 major to minor ratio that are broken up by a mitotic recombination event that occurs numerous times independently in the population.

**Fig 4.**
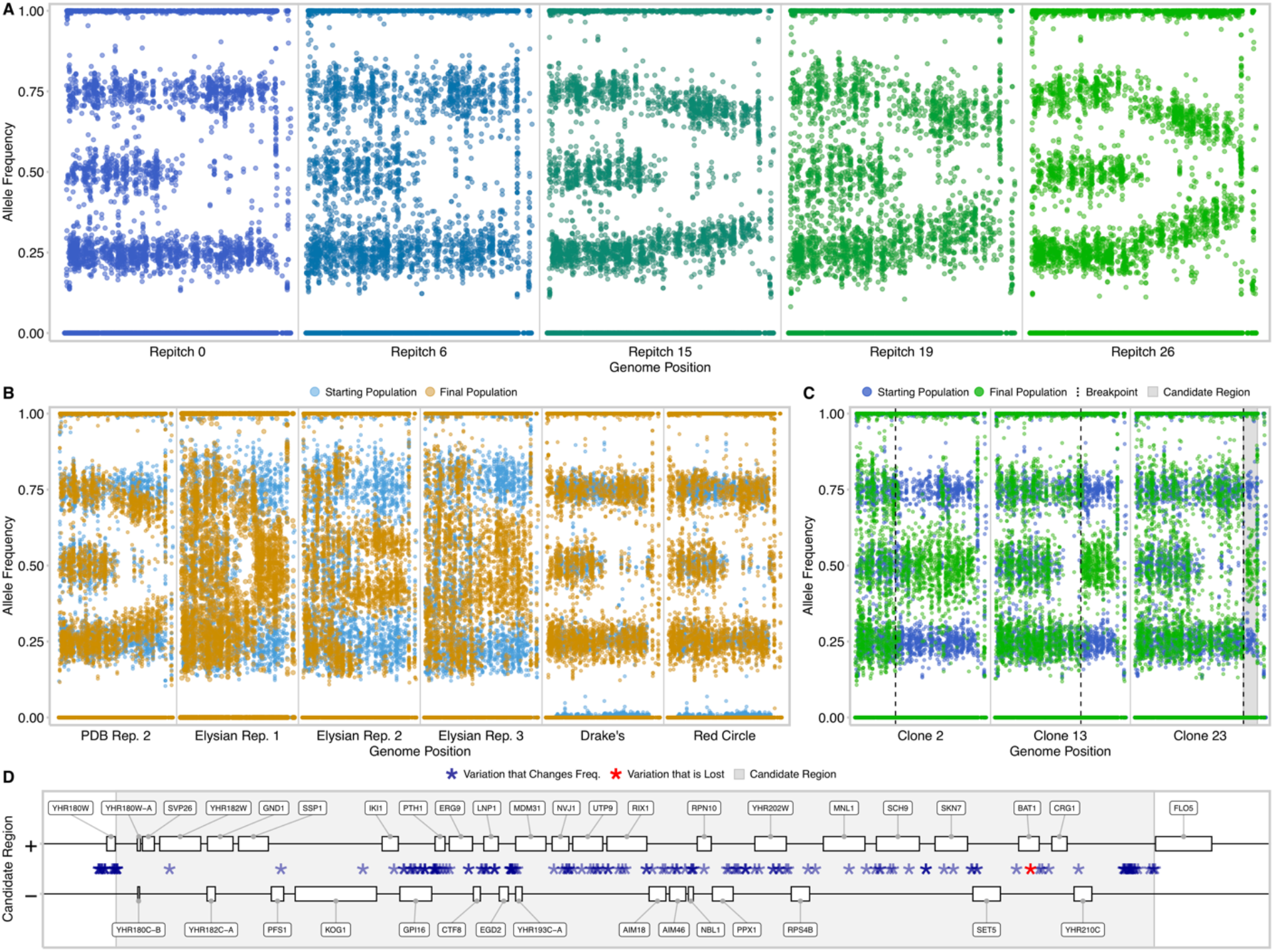
Mitotic recombination events spanning the same region recurrently mutated across multiple populations. (A) Allele frequency plots of serially-repitched populations from the first Postdoc Brewing Co. replicate showing an allele frequency change, appearing at the 15th repitch and reaching a 43.8% frequency in the population by the 26th repitch. (B) Replicate populations from Postdoc Brewing Co., Elysian Brewing Co., Drake’s Brewing Co., and Red Circle Brewing Co. showing the chromosome VIII allele frequency change. When the sample from Drake’s and Red Circle were compared to their ancestors, WLP001 and Cali Ale, it was found that the allele frequency change had previously occurred and is fixed in the strain. (C) Chromosome VIII of three representative clones from the 26th repitch from the first Postdoc Brewing Co. experiment showing the different breakpoints of the mitotic recombination event. The region used to detect lost variation is highlighted in grey. (D) The minimal region, as detected from clone 23 is displayed with all known ORFs and variation that is either lost or changes frequency as a result of the mitotic recombination.

Expanding our analysis to the other replicate populations, we observed that the second experiment from Postdoc Brewing Co. had a similar angled allele frequency change reaching 25.1% in the population, while all the Elysian Brewing Co. experiments showed sharp *de novo* breakpoints with different start locations (Fig 4B). Surprisingly, samples from breweries using the WLP001 strain from White Labs already had a starting fixed allele frequency change before entering the brewery. When we further investigated the rest of the American strains we found that the entire branch leading to WLP001 shared this breakpoint, indicating that it had occurred since its divergence from BRY-96 (Fig 2). Observing the same allele frequency change between multiple replicates at multiple breweries and independently within the American brewing yeasts, we concluded that this allele frequency change likely confers an adaptive benefit. From the Postdoc Brewing Co. replicate 1, for which we sequenced multiple intermediate samples, we estimated the selective benefit of the allele frequency change would be 5.70%, using a value of 3 generations per repitch.

We additionally observed chromosomes XII and XV experiencing convergent mitotic recombination events in 6 and 4 of the other populations respectively (Supp. Fig 4 and 5). After noting the mitotic recombination on chromosome XII in the other populations, we noticed that the first Postdoc Brewing Co. replicate likely had a similar event nearly fix in the population before it entered the brewery as one of the starting clones, Postdoc Brewing Co., timepoint 1 clone 1, did not have the allele frequency change. Using the clone that did not have the allele frequency change, we looked for any variation that experienced a LOH as a result of the mitotic recombination as this is the most likely source of an adaptive benefit for a mitotic recombination. However, through computational and manual inspection, we determined that no variation was lost as a result of the chromosome XII mitotic recombination (though other explanations are possible as well, such as allele copy number changes). Notably, the right arm of chromosome XII has been observed to have the highest amount of homozygosity among natural and industrial strains of yeast, potentially due to the presence of the rDNA locus on chromosome XII [3].

### Possible driver genes on chromosome VIII

Wanting to discover the basis of the selective benefit for the chromosome VIII mitotic recombination events, we also investigated whether any variation was eliminated as a result of this allele frequency change. We compared several clones from the first Postdoc Brewing Co. repitch experiment to identify the smallest candidate region in which the allele frequency change occurred, we then filtered for positions inside of the region where variation is lost (Fig 4C). As the SacCer3 reference genome does not capture the genome structure at the end of chromosome VIII and breaks down at the *FLO5* gene (see below), our analysis of lost variation spanned *YHR180W*to *FLO5* (Fig 4D). There was a single nonsynonymous mutation on one haplotype that was eliminated in every clone bearing a known allele frequency change and, notably, two clones without an allele frequency change (PDB26, clones 7 and 12). The mutation (an alanine-to-asparagine substitution) was found at position 234 in the gene *BAT1,* which encodes a mitochondrial branched-chain amino acid (BCAA) aminotransferase Bat1 that is critical in the metabolism of BCAA (valine, leucine, and isoleucine). Due to the importance of Bat1 for BCAA metabolism even beyond the context of this study, we analyzed the function of the A234D variant in a companion study (Jirasin Koonthongkaew et al., submitted [36]). Briefly, we discovered that in an otherwise isogenic background, the *BAT1* variant (*BAT1*^A234D^) leads to the same phenotype as a null allele in *BAT1.* Specifically, we found that both the null allele and the *BAT1*^A234D^ allele caused a growth defect in minimal medium, reduced levels of intracellular valine and leucine during the logarithmic and stationary phases, respectively, and produced more fusel alcohols.

Interestingly, we found that in the Elysian Brewing Co. experiments, *BAT1* did not contain the A234D allele, but the allele frequency change on chromosome VIII still occurred. Investigating these populations, we found no additional variation that was lost as a result of the mitotic recombination, leading us to suspect that there was additional gene content at the end of chromosome VIII that could be further driving the benefit of the mitotic recombination. However, the level of structural divergence between the SacCer3 reference genome and the beer strain was too great to be bridged using short read sequencing, especially due to the repetitive and paralogous nature of the flocculin gene. As there are no currently available long-read sequencing data for the American brewing strains, we generated our own using clones isolated from the first Postdoc Brewing Co. experiment.

### Oxford Nanopore Technology (ONT) based long-read analysis

Using a MinION sequencer, we generated ONT reads from a clone isolated from the first time point and three clones from the last time point (each with an allele frequency change). From these reads we generated individual assemblies from each of the sequencing runs and polished them for quality using both the ONT and Illumina reads (see methods). We found that multiple larger scale genome rearrangements had occurred at the telomeres of the beer strains. In particular, the right end of chromosome VIII had two separate rearrangements versus the SacCer3 reference that had occurred sometime in the ale brewing yeast past, one matching the left arm of chromosome I and the other the left end of chromosome IX. Furthermore we found that the sequence found at the left end of chromosome I had also transferred to the right arm of chromosome I. Based on previous literature we suspect that these two intragenomic recombination events have been referred to as *Lg-Flo1* (chimera between *FLO5* and *YAL065C* originally discovered in lager yeast) and *ILF1* (chimera between *FLO5* and *YIL169C)* [37]. Beyond the chimeric flocculins, we also discovered that additional gene content, extending to the telomere, was transferred. From the chromosome IX segment, *HXT12, IMA3, VTH1,* and *PAU14* were duplicated to chromosome VIII. From the chromosome I segment, *SEO1* and *PAU8* were duplicated.

Using alignments of the ONT reads back to polished assemblies, we established what variation and haplotypes were attached to which telomeric ends. Specifically, we found that the chromosome encoding the *BAT1*^A234D^ allele is connected to the fragment from chromosome IX. Additionally, the minor haplotype is connected to the fragment from chromosome I, while the remaining two chromosomes are connected to the content from chromosome IX. While these observations were confirmed using ONT reads from clone isolates from the Postdoc Brewing Co. experiment, we have further found that the copy number of the chromosome I fragment containing *SEO1* increases in both Postdoc Brewing Co. populations and the three Elysian Brewing Co. populations by the final timepoint, meaning that the copy number of *Lg-FLO1* and *SEO1* likely both increased in all populations that experienced a chromosome VIII mitotic recombination (Supp. Fig 6).

### Flocculation

As there was a change in the copy number of Lg-*FLO1*, we tested whether there were any changes in flocculation rate of clones isolated from the first versus the last time point of the Postdoc Brewing Co. first replicate experiment. We found that there are no substantial shifts between clones bearing the chromosome VIII allele frequency change and the clones that do not (Supp. Fig 7). However, because the experiments were conducted in small scale laboratory conditions in non-optimal media conditions to test for flocculation of beer brewing strains, more experimentation is required to conclusively eliminate the possibility that there are differences in flocculation speed or strength between the clones.

### Growth Phenotypes

Given the potential of an evolutionary benefit of the allele frequency change on chromosome VIII and the aneuploidy of chromosome V, we tested for any growth changes in brewers wort of clones bearing these mutations. Fitting growth curves of these yeasts with a linear model on the period of exponential growth, we analyzed whether there were any changes in the growth rate or lag time of the clones using a Kruskal-Wallis statistical test. Finding a difference in the growth rates (p-value = 3.718 x 10^-5^), we further probed for differences between clones using a Mann-Whitney U test. Consequentially, we found that the growth rate of the two clones with the chromosome VIII allele frequency change had a significantly increased growth rate versus the two clones isolated from the first timepoint (Fig 5).

**Fig 5.**
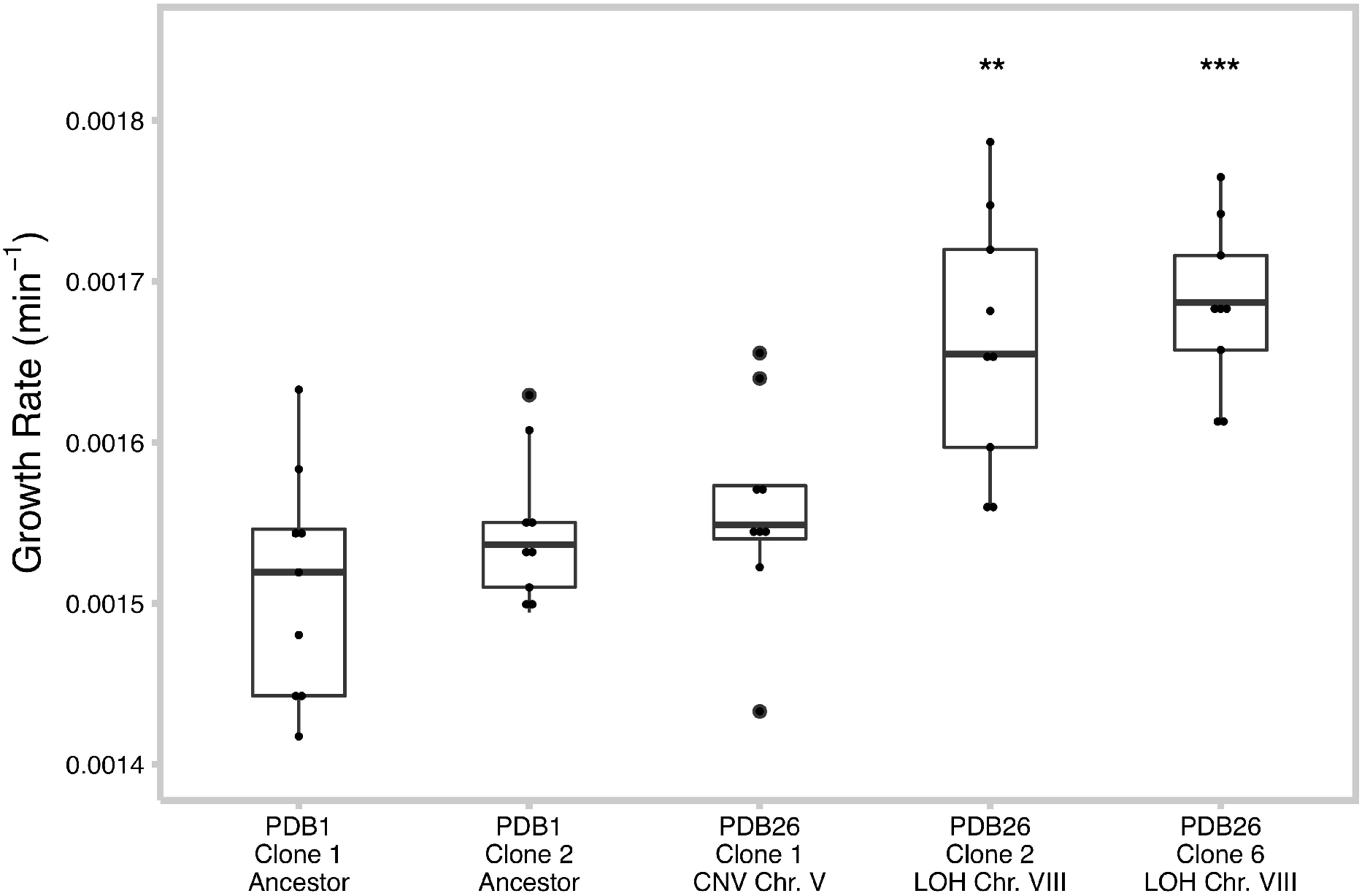
Clones bearing a Chr. VIII mitotic recombination grew significantly faster in brewers wort. Growth rates of clones isolated from the first Postdoc Brewing replicate were measured over 24 hours of growth in brewers wort using a plate reader and compared using a Kruskal-Wallis test (p-value = 3.718 x 10^-5^). Comparisons using a Mann-Whitney U test revealed significant differences between clones bearing the mitotic recombination on chromosome VIII (** p-value < 0.001; *** p-value < 0.0001).

### Changes in Sensory Profiles

More than anything, the phenotype that is most important for brewers is the taste of their beer. To assay for any changes in flavor, we brewed beer with the ancestral strain and two of the clones described above, and measured both the molecular profile of the beer and the sensory profile with a crowd-sourced panel from attendees at a Homebrewing Convention. Among the two fermentation replicates that we tested with chemical profiling, we found that there was an increased quantity of isobutanol isoamyl acetate and amyl alcohols in the clone lacking the *BAT1*^A234D^ allele (Table 3). Notably, our prior investigation of the A234D allele in a laboratory strain background conclusively found this same pattern, potentially indicating *BAT1* as the driver of the metabolite differences (Jirasin Koonthongkaew et al., submitted [36]).

**Table 3:** Sensory analysis of beers brewed with clones from first Postdoc Brewing replicate

| Sample Name | BAT1 Status | Ferment | Alcohol (%v/v) | Specific Gravity | Alcohol (%w/w) | pH Value | Acetaldehyde (ppm) | 1-Propanol (ppm) | Isobutanol (ppm) | Amyl Alcohols (ppm) | Iso-Amyl Acetate (ppm) | Diacetyl (ppb) | Total Diacetyl (ppb) | 2,3-Pentanedione (ppb) | Total 2,3-Pentanedione (ppb) |
| --- | --- | --- | --- | --- | --- | --- | --- | --- | --- | --- | --- | --- | --- | --- | --- |
| PDB1 Clone 1 | A234D | 1 | 5.1 | 1.007552 | 4 | 4.54 | 5.89 | 28.78 | 18.12 | 66.29 | 0.42 | 32.35 | 44.32 | <10 | <10 |
| PDB26 Clone 6 | + | 1 | 4.72 | 1.010501 | 3.69 | 4.65 | 6.64 | 17.82 | 8.26 | 39.82 | 0.28 | 74.83 | 116.86 | 11.96 | 18.83 |
| PDB26 Clone 1 | A234D | 1 | 5.1 | 1.007233 | 4 | 4.63 | 3.55 | 24.77 | 15.77 | 63.9 | 0.89 | 24.06 | 49.47 | <10 | <10 |
| PDB1 Clone 1 | A234D | 2 | 4.56 | 1.015936 | 3.55 | 4.12 | 3.03 | 24.8 | 43.28 | 93.42 | 0.18 | 73.55 | 89.61 | <10 | <10 |
| PDB26 Clone 6 | + | 2 | 4.5 | 1.016369 | 3.5 | 3.96 | 3.84 | 23.28 | 29.6 | 75.36 | 0.06 | 81.78 | 121.92 | <10 | 10.12 |
| Wort Control | N/A | 2 | 0 | 1.050506 | 0 | 4.41 | 0.65 | 5.28 | 0 | 0 | 0 | 37.83 | 41.66 | <10 | <10 |

Additionally, we observed from the specific gravity measurements that the fermentations with the clone that experienced the mitotic recombination on chromosome VIII (PDB26 clone 6) potentially did not go to completion when compared to the fermentations from the clones containing the *BAT1*^A234D^ allele (PDB1 clone 1, and PDB26 clone 1). While we don’t know if this is linked to this specific allele or another mutation that the clone has, this feature overwhelmed the sensory panel, who found the beer to be different in both its maltiness and sweetness (Supp. Table 6). As well, we found an increase in the production of diacetyl, total and otherwise, but we do not know definitively whether this was because of reduced ability to clean up the fermented product due to a fermentation delay. Further replicates and additional testing of clones genetically manipulated to alter the *BAT1* allele identity are warranted to conclusively test the impact of the chromosome VIII allele frequency change, with and without the *BAT1*^A234D^ allele, on beer characteristics.

## Discussion

Using whole-genome sequencing on yeast serially repitched across four breweries, seven populations, and three different strains, we observed the repeated occurrence of convergent mutations rising to high frequency in the populations. Notably, we observed multiple types of structural variation impacting chromosomes V, VIII, XII, and XV across multiple replicates. Through subsequent phenotyping of clones bearing some of these mutations, we have found a growth rate benefit in strains carrying the mitotic recombination event on chromosome VIII when grown in brewers wort. From these data we have concluded that these mutations are likely beneficial and selected for in the brewery, indicating that despite centuries of growth in the brewery, ale yeasts continue to show signatures of new adaptations.

Given the few number of convergent mutations, we sought to determine the driving force behind the potential adaptive benefit of the chromosome V copy number change and the chromosome VIII mitotic recombination. First, from clone and population sequencing, we were able to determine that the copy number change was not haplotype dependent, meaning that the benefit likely originated from a dosage balance with the rest of the genome. Second, we found that as a result of the mitotic recombination on chromosome VIII a point mutation in *BAT1* in the Postdoc Brewing Co. populations was recurrently lost. Creating the *BAT1*^A234D^ allele in a lab strain and comparing to an isogenic wild-type, we discovered that the mutation led to a number of phenotypes including a sensitivity to osmotic stress, reduced fermentation ability when grown in 20% glucose, and a growth defect in minimal media (Jirasin Koonthongkaew et al., submitted [36]). We similarly found that the ale strain clones bearing the *BAT1*^A234D^ allele have a growth defect, indicating its likely influence on the brewing yeast growth and the origin of at least part of the mitotic recombination benefit. However, the populations from Elysian Brewing Co. did not have the *BA T1*^A234D^ allele, indicating there is likely to be additional adaptive consequences from the mitotic recombination event. Therefore, we applied long-read sequencing and *de novo* assembly on the clones from the first Postdoc Brewing Co. replicate to resolve the structure of the telomeric regions. As a result we found that the mitotic recombination led to a copy number increase of *Lg-FLO1* and *SEO1* from the minor haplotype and reduction of *ILF1, HXT12, IMA3, VTH1,* and *PAU14* from the major haplotype. Given the alteration in copy number of multiple flocculation associated genes, we tested several clones with and without the mitotic recombination and found no strong difference in the rate that they settled. Considering the lack of an obvious flocculation difference, we suspect that either our methods to measure flocculation were not sufficient to detect a significant change or that the benefit was derived from either the loss of the major haplotype gene copies, or the gain of *SEO1,* whose function is presumed to be related to nitrogen uptake [38]. Notably, there are six copies of *SEO1* in the genome; however, this might be coincidental due to its linkage to flocculin-associated gene sequences.

Considering the long history of beer brewing, one might presume that the yeast specialized in malt fermentation would already be pre-adapted to the brewery environment. Especially given that repitching is not a new phenomenon with Louis Pasteur commenting in 1876 on the practice of passaging yeast within and between breweries:

> [T]he wort is never left to ferment spontaneously, the fermentation being invariably produced by the addition of yeast formed on the spot in a preceding operation, or procured from some other working brewery, which, again, had at some time been supplied from a third brewery, which itself had derived it from another, and so on, as far back as the oldest brewery that can be imagined.… [T]he interchange of yeasts amongst breweries is a time-honoured custom, which has been observed in all countries at all periods, as far back as we can trace the history of brewing. ([39], p. 186)

However, within our experiments we see multiple mutations overtaking the population in a relatively short period of time. The most parsimonious answer is that the brewing process has somehow changed in a way that creates new selective pressures, allowing for novel, highly beneficial mutations to evolve. One possibility is that the breweries which we partnered with utilize styles that the ‘Chico’ yeasts had not been extensively exposed to. Specifically, in the United States, styles of beer that are high in both final alcohol and hop content have become popular and are utilized extensively by Postdoc Brewing Co., Drake’s Brewing Co. and Elysian Brewing Co. Seeing as high hop and alcohol content can be stressors, we hypothesize that this could be one of the contributors to new adaptations. Further experimentation using defined media, varied in both hop and sugar content, will be able to test this hypothesis. Another possibility is that industry has shifted in the last several decades towards the use of pure clonal strains and propagation companies for yeast maintenance versus keeping yeast at scale in the brewery constantly through continual reuse. Often, to create a stock of a brewing strain the population is bottlenecked down to a very small size, making a single representation of the population. Through this process it is likely that mutations that aren’t representative of the population and are potentially deleterious become fixed. Further exacerbating the problem, this process repeats every time a new propagation company creates their own version of a strain. When the strain is then grown to a massive population size in stressful brewing conditions, there is then selection for *de novo* reversions of the deleterious mutations. Within our brewing experiments, we have seen two examples that potentially fit this explanation. First, the copy number of chromosome V returned to a euploid copy number, which was the hypothesized ancestral state. Second, the nonsynonymous mutation in *BAT1* was reverted by mitotic recombination. While both of these hypotheses would potentially explain the source of new adaptations, further work is required to rigorously test their veracity.

Another interesting complication for creating a single strain representation of a population was the occurrence of multiple lineages in the Postdoc Brewing Co. experiments that had the same or similar mutations within the population. This phenomenon, called clonal interference, occurs when a new beneficial mutation is unable to completely overtake the population before another beneficial mutation occurs. This creates competition between the new and old beneficial mutation, preventing a single lineage from taking over the population. Typically the parameters that are thought to control the degree of clonal interference during adaptive evolution are the mutation rate of beneficial mutations, the selective benefit of those mutations, and the population size [40]. As mentioned above, breweries have an immense population size, creating an ideal environment for clonal interference. However, it is unclear why in the non-Postdoc Brewing Co. populations we did not see the same degree of clonal interference. We suspect that the euploid nature of chromosome V of BRY-96 and the preexisting mitotic recombination in WLP001 may have allowed for a different population dynamic that led to a single lineage dominating the population. The other possibility is that the beneficial mutations observed in the non-Postdoc Brewing Co. populations occurred earlier in their outgrowth, leading to a single lineage dominating the population.

Matching with this hypothesis, we found that on multiple instances the yeast entering the brewery already had undergone some amount of genetic divergence from the stock’s genotype. This is likely due to the number of generations required for a stock of yeast to be grown to a population size needed by professional brewers. For example, given a 20 hectoliter batch of wort, the recommendation by White Labs, a prominent propagation company, is to add 2.4 x 10^13^ cells of yeast. The absolute minimum number of generations required to reach this number of yeast from a single cell, assuming a doubling per generation with no death, is 44.4 generations of yeast growth, which is almost certainly an underestimate. The number of generations occurring in the brewery, assuming 3 generations per beer fermentation, is 45 generations for 15 serial repitches and 78 for 26 serial repitches, meaning the growth period at the propagation company constitutes anywhere from half to a quarter of the yeasts growth in our experiments. As well, because mutations enter the population at a proportion of one over the total population size, beneficial mutations that occur earlier in the outgrowth have a higher probability of reaching a high frequency. Given the number of yeast cells needed by a brewer, it is likely inevitable that some amount of detectable evolution will occur prior to a pitch even entering the brewery.

Typically, the professional brewer wants to know how long they can reuse their yeast before they will start to notice considerable difference in the characteristics of the yeast or beer. Perhaps the most accurate but somewhat unsatisfying response is that it depends on a number of factors. Specifically, the timing might be different given the spectrum of adaptive mutations that a particular strain has access to, the individual mutation rate of that isolate, and the number of generations that the population was grown at the propagation company. Even given replicates using the same strain, there is an element of stochastic mutation that can potentially drastically change how a brewery population evolves. Looking to the future and at methods to have serially repitched populations with fewer impactful mutations may begin with sequencing more populations and finding isolates that are better preadapted to the modern brewery. However, this strategy assumes these mutations do not have undesirable tradeoffs on other aspects of performance, such as on flavor profile. As well, if the possible spectrum of adaptive mutations is determined for a given strain it may be possible for a commercial service to track the frequency of these mutations over time and identify when they start impacting the beer. We note that due to the variability in beer styles employed during most of the time courses analyzed here, we were unable to rigorously track changes in fermentation characteristics and/or beer quality that may have happened in tandem with the rise of these mutations.

In conclusion, we observed multiple independent brewing yeast populations with high-frequency structural mutations that likely contributed to a change in growth characteristics. Discovering the likely adaptive benefit of mitotic recombination events in the brewery raises the possibility that historical ale brewing yeast adaptation was due in part to these kinds of structural mutations. Notably, the ale yeasts are thought to have originated from an admixture event which introduced intragenomic variation into the ancestor of the modern brewing strains [8]. Potentially, ale yeasts have the capacity to adapt to new conditions using mitotic recombination on existing variation to eliminate or fix deleterious or adaptive alleles respectively. Such events have been observed within the lager brewing yeasts wherein similar patterns of structural variation have been linked to phenotypic outcomes [41,42]. Furthermore, multiple mitotic recombination events were shown to lead to lead to changes in both sugar utilization and flocculation when *de novo* hybrids between *S. cerevisiae* and *S. eubayanus* were evolved in simulated brewing conditions [20]. Given the prevalence of structural mutations in the history of the genome of brewing yeasts, and their link to adaptive phenotypic outcomes, further investigation into the consequences of this variation will likely provide additional insights into their domestication.

## Materials and Methods

### Evolution in the Brewery

Depending on the brewery, yeast cells were ordered from Wyeast, Imperial Yeast, White Labs, Escarpment Laboratories, or an internal propagation service (in the case of Elysian Brewing Co.). For some of the experiments, starting samples were collected either from the shipment or from the first beer brewed with the yeast. Otherwise these yeast cells were commonly grown for several generations in low density wort, then transferred into a cycle of several ale beer recipes ranging from barley wine to double IPAs. The precise recipe and conditions are proprietary for some of the breweries, however Postdoc Brewing Co. has provided the style in which the yeast were passaged through (Supp. Table 1). For the Postdoc Brewing Co. samples, they were collected from the middle of the flocculated yeast after the runnings of hop and protein particulate was disposed of. Once samples were collected, they were stored at 4°C in a sterile, airtight container until transfer to the laboratory was possible. Upon arrival in the laboratory, the samples were thoroughly mixed and 1 mL was transferred to a 25% glycerol stock that was subsequently frozen at −70°C.

### Short-read genome sequencing

Populations of yeast cells, previously stored in 25% glycerol at −70°C were transferred to deionized water (diH_2_O) and measured for cell density using a hemocytometer. Based on cell density counts in the diH_2_O, the cell suspensions were diluted and plated to collect approximately 1,000 independent yeast colonies, grown for 4 days on yeast extract peptone dextrose (YEPD) plates with 2% glucose and 1.7% agar at room temperature. These plates were scraped for cells with a sterile glass rod, concentrated by centrifugation, and washed in diH_2_O. DNA was then extracted from the cell pellets using a modified Hoffman-Winston preparation [43].

Single clone isolates were generated from a population glycerol stock. In short, the brewery populations were streaked onto a YEPD plate and grown at room temperature. A single colony was isolated and grown overnight in 5 mL of YEPD liquid medium with rotation. A portion of the overnight culture was stored in a 25% glycerol stock for archiving and subsequent analysis. The remaining cells were concentrated, washed with diH_2_O and had their DNA extracted with a modified Hoffman-Winston preparation [43]. Clones 13 through 23 were selected for sequencing based on their likelihood for bearing a chromosome VIII allele frequency change from genotyping using PCR and Sanger sequencing for a SNP frequency within a variable region on the end of the chromosome.

After measuring the concentration of DNA using a Qubit Fluorometer (Thermo Fisher Scientific), dual-indexed Illumina libraries were generated using a Nextera sample preparation kit (Illumina, Inc.) with 50 ng of input DNA. The genomic libraries were sequenced using 150-bp paired end sequencing on an Illumina NextSeq 500 using the manufacturer’s recommended protocols.

### Whole genome analysis

The Illumina reads were demultiplexed using bcl2fastq with default parameters. The reads were then aligned to the SacCer3 reference genome (R64-2-1) using BWA-mem (version 0.7.15) [44]. The alignments, after being sorted and indexed with SAMtools [45] (version 1.9) were marked for duplicates using Picard Tools (version 2.6.0). When libraries were sequenced on multiple lanes or runs, the alignments were combined using SAMtools. Afterwards, the alignments had their InDels realigned using GATK (version 3.7).

Short mutations such as SNPs and InDels were then called using three separate variant calling software packages. First, BCFtools call using modified input parameters was used. Second, FreeBayes (version 1.0.2-6-g3ce827d) [46] using input parameters (–pooled-discrete – pooled-continuous-report-genotype-likelihood-max –allelebalance-priors-off –min-alternate-fraction 0.1) were used to call both SNPs and InDels. Finally, in a paired mode with the sample’s ancestor, LoFreq was used to call SNPs [47]. For all of the variant callers, BEDtools was used to filter the variants called for a sample versus its ancestor [48]. Each variant file was subsequently filtered using standard parameters that are listed in Supp. Table 7. The three variant files were then filtered to exclude overlaps of the same variant and combined into one file using a custom script. Afterwards, the annotation and impact of the variants were determined using a script previously published in [49]. Finally, each variant that passed all filters was manually checked for its authenticity in the Integrative Genomics Viewer (IGV) [50]. When variant calls from BCFtools call exceeded 300 variants, these files were ignored, as they were found to contain primarily false-positives through manual inspection and comparisons with other software.

As noted earlier, there are multiple haplotypes containing varying degrees of shared variation between homologous chromosomes. To quantity and observe the degree that this variation has been altered through mitotic recombination, allele frequency was calculated and plotted for all genomic coordinates. Briefly, from the previously generated alignments, variant calls were generated using the GATK (version 3.7) HaplotypeCaller. These variant calls were passed to GATK VariantToTable and modified using an in-house java script into a per base allele ratio between a reference and alternate allele. Subsequently, the allele frequency was plotted using an R script with ggplot2. Changes in the ratios between haplotypes were visually determined through inspection of these plots. Precise values on the proportion of the allele frequency change of chromosome VIII in the population were generated using an average of the change in allele frequency of a set of SNPs that were highly representative of the mitotic recombination events in the clones at the end of chromosome VIII. These values were then used to calculate the selective benefit of the chromosome VIII allele frequency.

Using the alignments listed above, the copy number of the genome was determined and plotted using an in-house script. Briefly, the total genome coverage was calculated using GATK (version 2.6.5) DepthOfCoverage. Next, the per window average coverage across the genome was calculated using the command-line tools version of IGVtools. These files were combined using a python script to generate a normalized coverage measure. As many of the samples included a ‘wavy’ coverage in which the coverage varied across the genome in an inconsistent and seemingly random pattern, the per ORF coverage was unable to be accurately determined. In the cases that the coverage was too ‘wavy’ to accurately determine the coverage, the allele frequency plots that are described above were used to determine the copy number as the per allele coverage remained unchanged by the ‘wavy’ sequencing artifact.

### Phylogenomic analysis

In order to properly understand the diversity and previous evolutionary history of the American brewing strains, all publicly available brewing strains whole genome sequencing were processed into a phylogenetic representation. Capturing the most amount of American diversity possible, some strains that had not previously been sequenced, but suspected to be part of the American yeast group (due to tips from professional and amateur brewers) were ordered and kindly donated from a variety of yeast propagation companies. As described above, the strains had their DNA extracted and sequenced using the paired-end Illumina technology. All of the sequencing reads were aligned using a similar strategy as previously described above with slight modifications, and called for variants in the GVCF mode using GATK (version 4.1.1.0) HaplotypeCaller on regions of high confidence (excluding the first and last 50 kb of each chromosome to avoid poorly assembled telomeric sequences). Individual variant calls were collected and jointly genotyped using GATK GenomicsDBImport and GenotypeGVCFs and filtered with GATK (version 4.1.3.0). Removing the influence of ancestral variation lost by mitotic recombination, the SNPs from the sample excluding BE051 were then filtered by SNPs called from BE051 using BEDtools [48]. The SNP calls were then converted into two concatenated fasta files wherein the first fasta was the SacCer3 reference genome with the reference allele as listed if the strain was either heterozygous or homozygous for the reference allele. The second fasta also contained the SacCer3 allele unless a heterozygous or homozygous variant position was detected in which case the alternate allele was outputted. This task was done using BCFtools. The concatenated fastas from all of the American brewing strains were passed to IQTree2 to generate a maximum-likelihood tree using GTR_4_ + gamma model [51]. The tree was then modified for aesthetics and annotation using iTOL [52].

Comparisons between the ‘Chico’ yeasts and BRY-96 for determination of the ancestry of the ‘Chico’ yeasts was done using the aforementioned SNP calls. First, the union of the SNPs called in WLP001 and Wyeast 1056 was generated using BEDtools. Second, the mutations unique to BRY-96 when compared with that union were generated. Finally, the remaining SNPs from BRY-96 were manually inspected for veracity using IGV.

### Flocculation

The rate of flocculation was quantitatively measured similar to previously reported [53]. Briefly, yeast of the appropriate genotype, plated on a 2% YPD plate were grown from a single colony in 5 mL of 2% YPD liquid medium for 72 hours at 30°C with rotation. The yeast cultures were then vortexed for a minimum of 5 seconds to ensure complete resuspension. Photographs were then taken of the yeast after 60 minutes while the culture tubes remained undisturbed. Afterwards, using a semi-automated script written for ImageJ [54], the images were converted to black and white, and the plot profiles of the culture tube’s grey intensity were collected from the bottom of the tube to the meniscus. Next, to determine the degree of settling in an unbiased manner, an automated script written in python was used to find the point in the culture that the intensity reached half of the maximum grey value. The point at which the yeast had flocculated to in the culture tube after 60 minutes was used to create a ratio based on the total length of the culture. Three measurements were taken per culture and the average of these measurements was reported. Two biological replicates were conducted from independent colonies.

### Brewers wort media

The brewers wort media, utilized for the growth phenotyping and fermentation analysis was made as previously mentioned in [55] with slight modification. Briefly, 320 grams of amber liquid malt extract from Breiss Malt and Ingredients Co were mixed with 1.5 liters of distilled water and boiled for an hour. Fifteen minutes before the boil finished, 0.2 gram of the Wyeast Beer Nutrient Blend was added to the mixture according to the manufacture’s guidelines. After the wort had been chilled to a workable temperature, the specific gravity was measured using a hydrometer (and the value read was corrected based on the temperature), and the media was passed through fresh Melitta filters to remove any large coagulants. Next, the media was passed through a 0.45 micron filter (Nalgene 500mL Rapid-Flow Bottle Top Filters) to completely sterilize the media. The specific gravity of all batches used herein were found to be the same value of 1.050.

### Growth phenotypes

The growth characteristic of clones isolated from the first replicate population from Postdoc Brewing Co. was measured. First, single yeast colonies from a 2% YPD plate were grown in wort medium for 48 hours with rotation. Next the optical density of the cultures was measured at 600 nm (OD600). Each culture was then diluted in an appropriate amount of wort to reach a final OD600 of 0.1. The back-diluted cultures were further transferred to a 96-well plate at a volume of 200 microliters per well. Using a Biotek Synergy H1 plate reader, the OD600 of the 96-well plate was measured every 15 minutes for 24-48 hours while shaking in a double orbital pattern at room temperature. Utilizing a script written in the R programming language, the growth data from the plate reader were analyzed using the growthrates package. Employing the growthrates implementation of fitting linear models to the exponential growth period outlined in [56], we extracted the maximum growth rate of the clones and the length of the lag growth period. To determine whether there was a difference in the growth rates between clones, we first conducted a Kruskal Wallis rank sum test. Further testing of differences between clone growth rates was done using Mann-Whitney tests.

### Sensory profiling

The isolated clones from the first Postdoc Brewing replicate were tested for differences in the production of flavor compounds and the effect these compounds had on the beer sensory profile. First, two separate beer batches (beer batches 1 and 2) were generated from fermentations carried out either at Postdoc Brewing (using an all grain pale ale recipe) or in the laboratory (using the malt extract wort mentioned earlier). The yeast that fermented the beer were grown in the laboratory from single colonies to the desired cell count in 2% YEPD liquid medium with shaking. For the first and second beer batch, the yeast were concentrated with centrifugation and pitched into the wort at a rate of 3.5 x 10^5^ and 1.0 x 10^6^ cells per degree of plato respectively.

Second, both beer batches were submitted to White Laboratories for analytical services including gas chromatography measurements of a number of flavor compounds and measurements of alcohol percentage and specific gravity. Next, the beers from batch 1 were submitted to an untrained judging panel (n=95) at the Homebrew Con 2018, who used a standardized beer scoresheet from the Beer Judge Certification Program to analyze the profile of the beer. The identity of the beers were kept masked from the participants while they filled out their analysis. Afterwards, the scoresheets were aggregated and analyzed for differences using a Kruskal Wallis rank sum statistical test.

### Oxford nanopore sequencing

Yeast cell cultures were grown overnight at 30°C in 20 mL of YPD medium to early stationary phase before cells were harvested by centrifugation. Total genomic DNA was then extracted using the QIAGEN Genomic-tip 100/G according to the manufacturer’s instructions. The extracted DNA was barcoded using the EXP-NBD104 native barcoding kit (Oxford Nanopore Technologies) and the concentration of the barcoded DNA was measured with a Qubit 1.0 fluorometer (Thermo Fisher Scientific). The barcoded DNA samples were pooled with an equal concentration for each strain. Using the SQK-LSK109 ligation sequencing kit (Oxford Nanopore Technologies), the adapters were ligated on the barcoded DNA. Finally, the sequencing mix was added to the R9.3 flowcell for a 48 hour run.

### Assembly generation and polishing

The ONT reads were demultiplexed using Guppy with default parameters. The adapters on the raw reads were removed using Porechop. Afterwards, each sample was independently run through SMARTdenovo with default parameters to generate a draft genome assembly. To improve the quality of the assembly, the draft sequences were first run through racon then medaka. Next they were refined using pilon and the Illumina reads previously generated for the four clones sequenced on the MinION. The identity of the contigs was determined through pairwise alignment of the contigs (masked with RepeatMasker) to the SacCer3 reference genome using Minimap2 and plotted using an R package called DotPlotly. Confirmation of the contig identity, and the inferred identity of the ancestor was done using a combination of Minimap2 alignments of the SacCer3 reference ORFs, SacCer3 reference sequence, ONT reads, and Illumina reads, all visualized in IGV.

## Data availability

All whole-genome sequencing data was uploaded to https://www.ncbi.nlm.nih.gov/sra with the BioProject accession number: PRJNA641752.

## Acknowledgments

Massive thanks to Professor Mary Kuhner for advice on best practices for complicated phylogenetic inference. Sequencing analysis consultation was graciously provided by Mitchell Vollger and Dr. Glennis Logsdon. Thanks to Jess Caudill at Imperial Yeast for insight to the yeast propagation industry. Thanks to the attendees of Homebrew Con 2018 who participated in the sensory analysis. Thanks to Karen Fortmann and White Labs for metabolite profiling. This research was only made possible by the generous contribution of numerous unlisted, but greatly appreciated brewers, historians, and researchers. This work was funded by grant 1516330 from the National Science Foundation. This material is based in part upon work supported by the National Science Foundation under Cooperative Agreement No. DBI-0939454. Any opinions, findings, and conclusions or recommendations expressed in this material are those of the author(s) and do not necessarily reflect the views of the National Science Foundation. The research of MJD was supported in part by a Faculty Scholar grant from the Howard Hughes Medical Institute. CRLL was supported by T32 HG000035 from the National Human Genome Research Institute. JS was supported by the Agence Nationale de la Recherche (GM101091-01) and the European Research Council (ERC Consolidator grant 772505). AT was supported by a grant from the Ministère de l’Enseignement Supérieur et de la Recherche. YT was supported by a Grant-in-Aid for Research Activity start-up (19K21144) from the Japan Society for the Promotion of Science (JSPS)

## Supplementary Figures

**Supplementary Fig 1.**
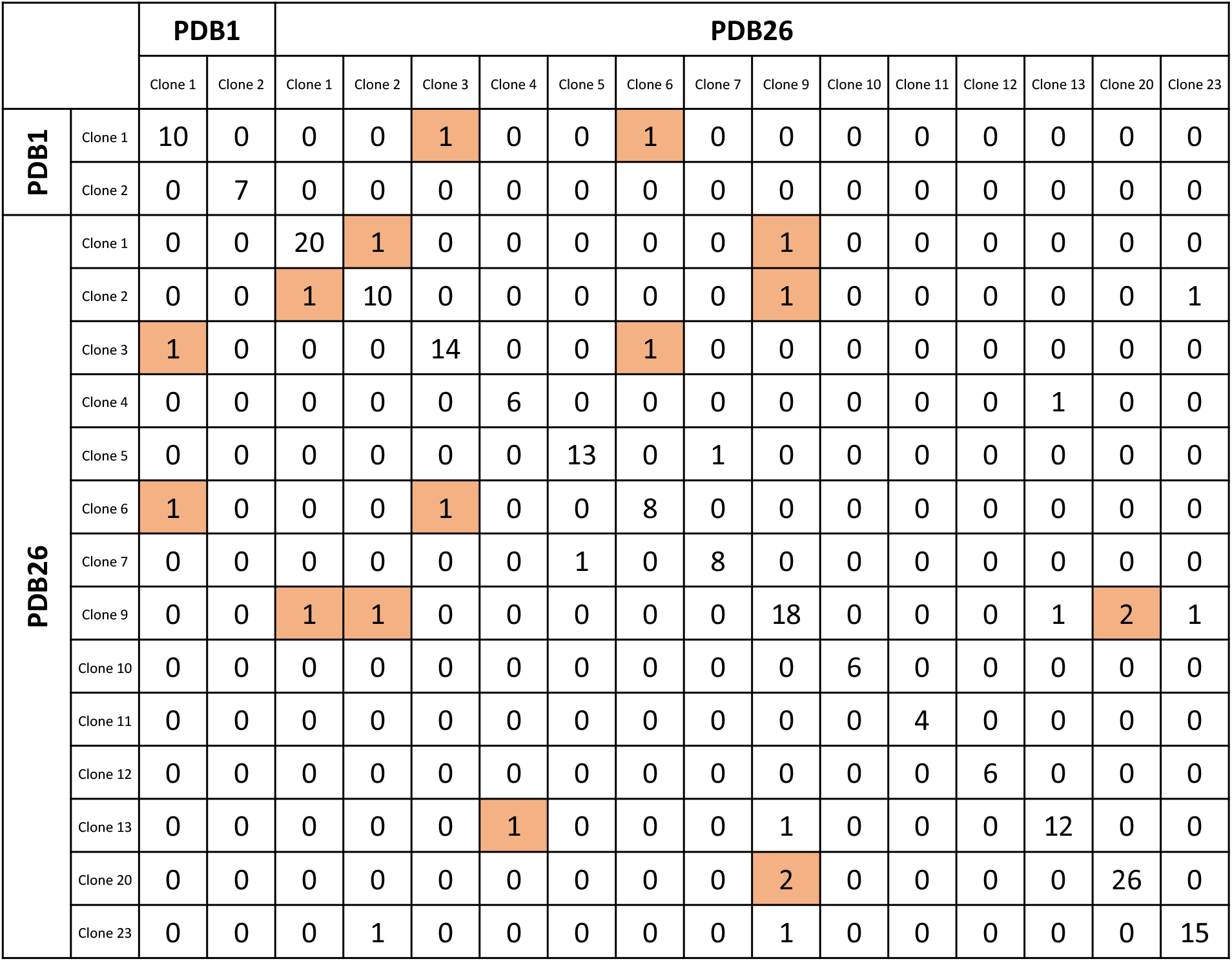
The number of mutations shared between clones isolated from the first Postdoc Brewing replicate experiment. The orange highlight indicates mutations that were also observed in the first or last timepoint populations from the first Postdoc Brewing replicate.

**Supplementary Fig 2.**
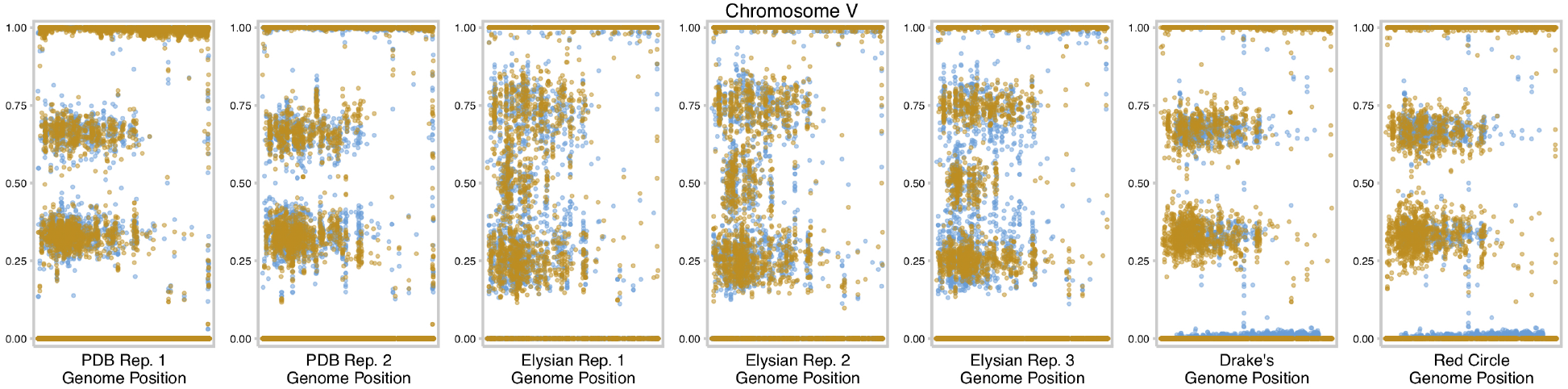
Allele frequency of chromosome V with the first timepoint colored in blue and the final timepoint in orange.

**Supplementary Fig 3.**
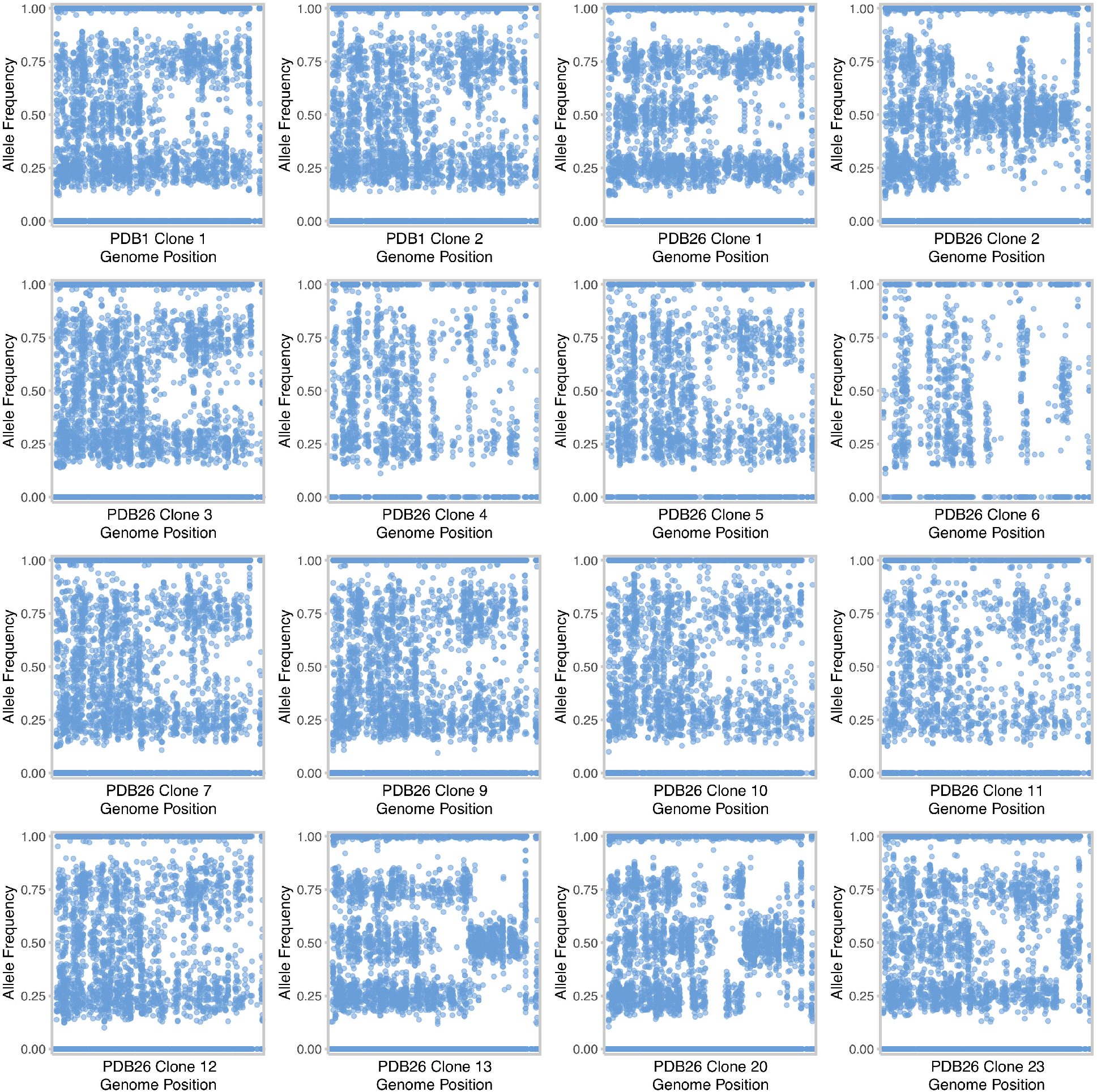
Allele frequency of chromosome VIII for every clone isolated from the first Postdoc Brewing Co. replicate.

**Supplementary Fig 4.**
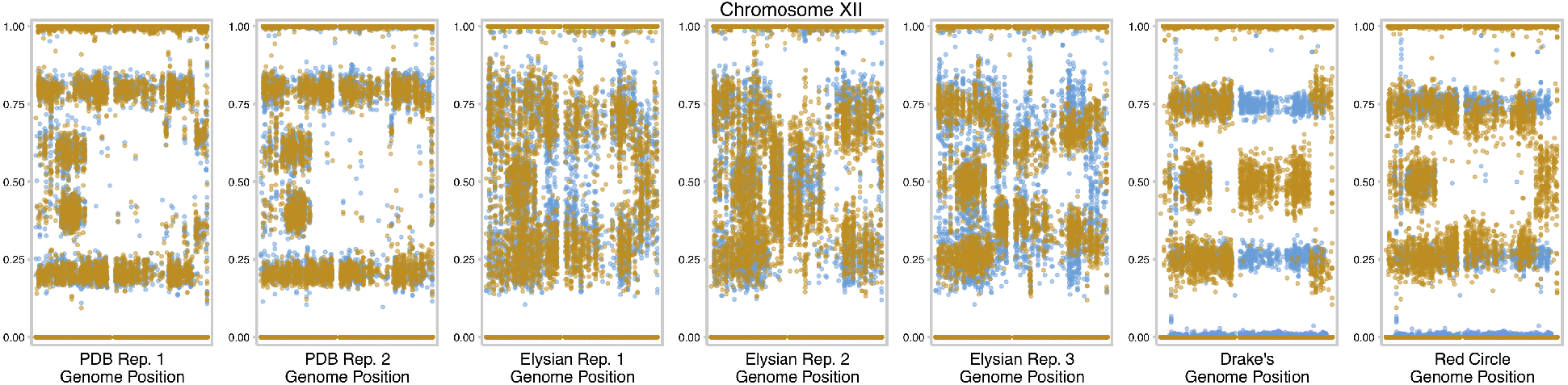
Allele frequency of chromosome XII with the first timepoint colored in blue and the final timepoint in orange.

**Supplementary Fig 5.**
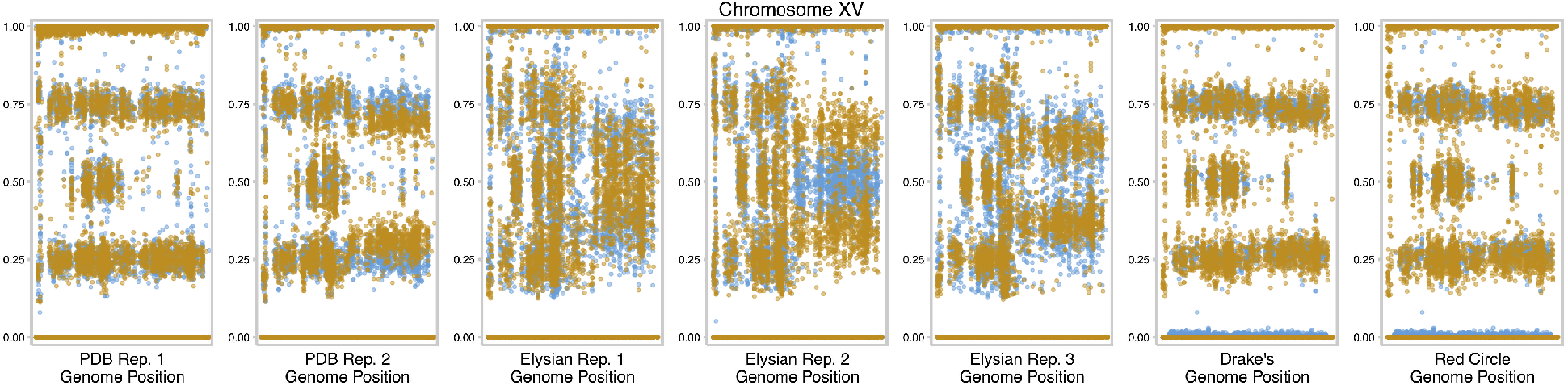
Allele frequency of chromosome XV with the first timepoint colored in blue and the final timepoint in orange.

**Supplementary Fig 6.**
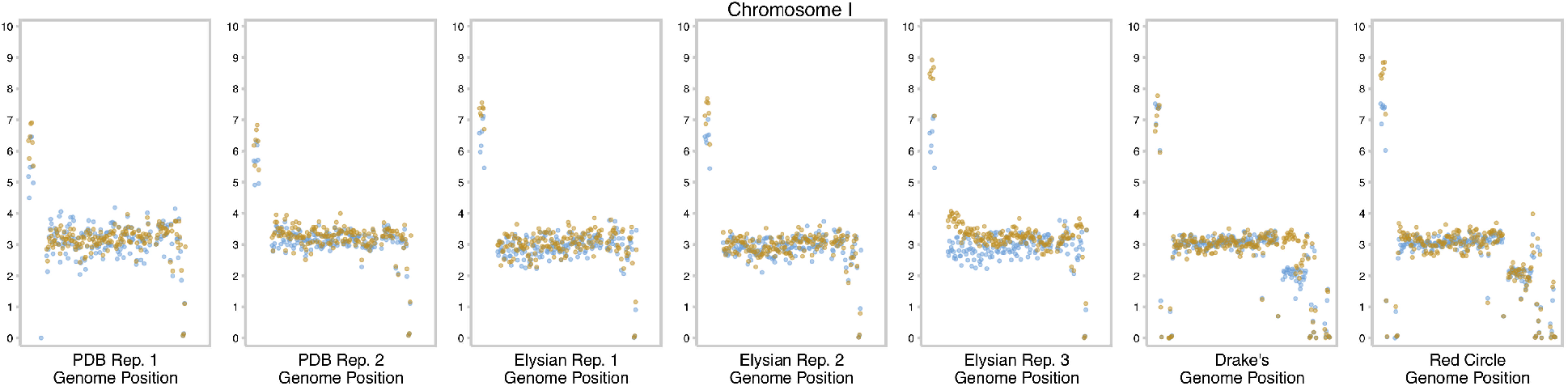
Copy number of chromosome I shown as 1000-bp sliding windows with the first timepoint colored in blue and the final timepoint in orange.

**Supplementary Fig 7.**
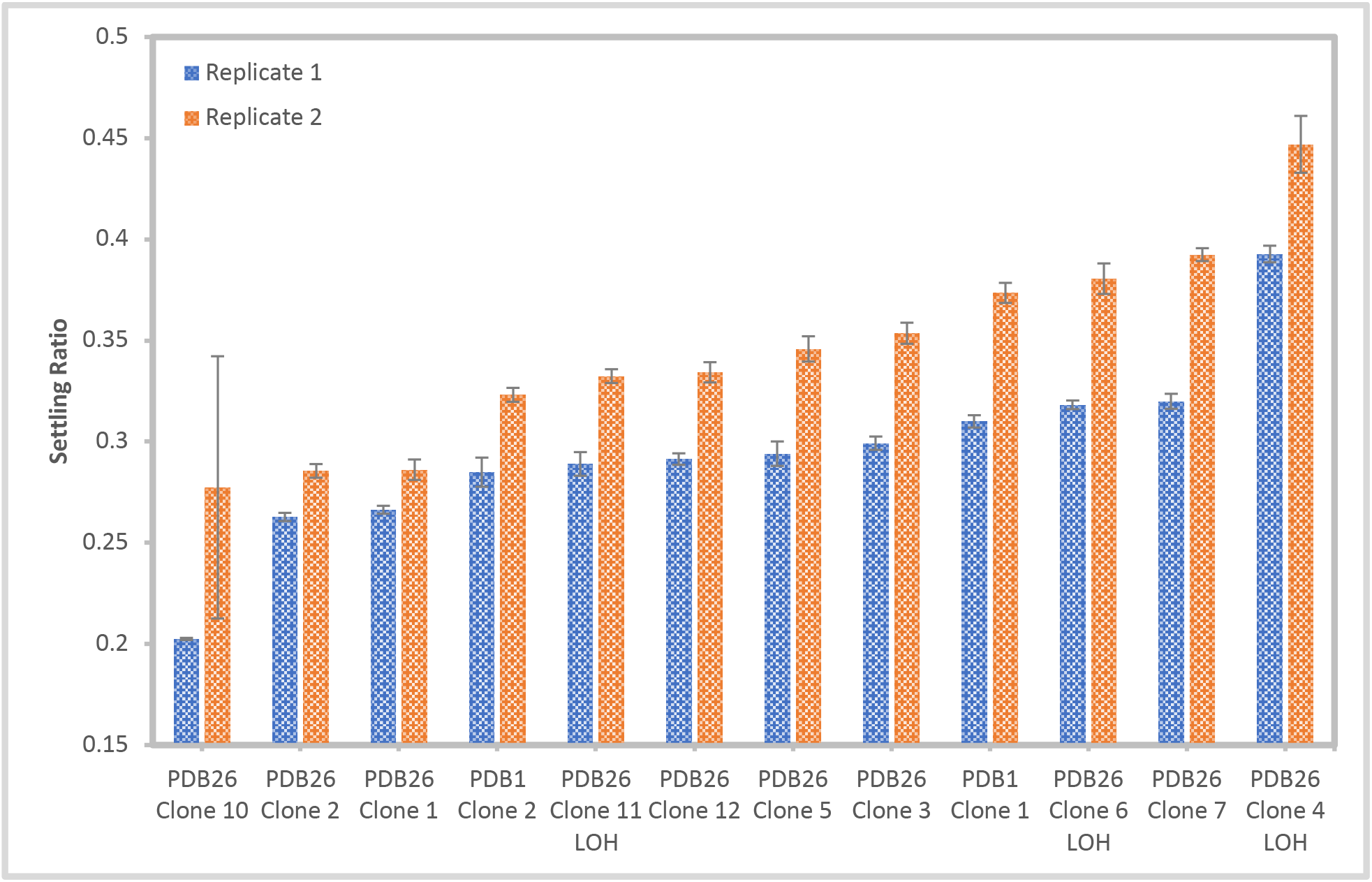
Settling ratio of clones isolated from the first Postdoc Brewing Co. replicate.

